# The Cannabis Multi-Omics Draft Map Project

**DOI:** 10.1101/753400

**Authors:** Conor Jenkins, Ben Orsburn

## Abstract

Recently we have seen a relaxation of the historic restrictions on the use and subsequent research on the *Cannabis* plants, generally classified as *Cannabis sativa* and *Cannabis indica*. What research has been performed to date has centered on chemical analysis of plant flower products, namely cannabinoids and various terpenes that directly contribute to phenotypic characteristics of the female flowers. In addition, we have seen many groups recently completing genetic profiles of various plants of commercial value. To date, no comprehensive attempt has been made to profile the proteomes of these plants. We report herein our progress on constructing a comprehensive draft map of the *Cannabis* proteome. To date we have identified over 17,000 potential protein sequences. Unfortunately, no annotated genome of *Cannabis* plants currently exists. We present a method by which “next generation” DNA sequencing output and shotgun proteomics data can be combined to produce annotated FASTA files, bypassing the need for annotated genetic information altogether in traditional proteomics workflows. The resulting material represents the first comprehensive annotated FASTA for any *Cannabis* plant. Using this annotated database as reference we can refine our protein identifications, resulting in the confident identification of 13,000 proteins with putative function. Furthermore, we demonstrate that post-translational modifications play an important role in the proteomes of *Cannabis* flower, particularly lysine acetylation and protein glycosylation. To facilitate the evolution of analytical investigations into these plant materials, we have created a portal to host resources we have developed from proteomic and metabolomic analysis of *Cannabis* plant material as well as our results integrating these resources. All data for this project is available to view or download at www.CannabisDraftMap.Org

## Introduction

Proteomics is a science dedicated to the creation of comprehensive quantitative snapshots of all the proteins produced by an individual organism, tissue or cell.^1^ The term was coined in the 1990s during the efforts to sequence the first complete human genomes.^2^ While the technology was in place to complete the human genome draft in 2003, the first two drafts of the human proteome were not completed by teams led by Johns Hopkins and CIPSM researchers until 2014. These two separate and ambitious projects were the composite of thousands of hours of instrument run time utilizing the most sophisticated hardware available at that time.^3,4^ Recent advances in mass spectrometry technology now permit the completion of proteome profiles in more practical time. Single celled organisms have been “fully” sequenced in less than an hour, and by use of multi-dimensional chromatography, relatively high coverage human proteomes have been completed in only a few days.^5–7^ While much can be learned by sequencing DNA and RNA in a cell, quantifying and sequencing the proteome has distinct advantages as proteins perform physical and enzymatic activities in the cell and are therefore more directly linked to phenotype.^8^ RNA sequencing may correctly predict the presence and relative abundance of proteins, but proteins inactivated by chemical modifications may make predictions of function from RNA abundance data wholly inaccurate. Furthermore, many proteins are altered by chemical post-translational modifications such as phosphorylation and acetylation which may completely change the protein function by serving as on/off switches for motion or metabolism.^9,10^ Protein modifications are directly involved in nearly every known disease and these modifications are impossible to identify with any current DNA/RNA sequencing technology.^11^

In North America we have recently witnessed an alleviation of restrictions on the use and subsequent research of plants belonging to the *Cannabis* genus. To date, relatively little work has been performed on these plants in any regard and no comprehensive study of the proteome has ever been attempted. In a study published in 2004, Reharjo *et al*., described a differential proteomics approach for the studying of *Cannabis sativa* plant tissues. Differential analysis was performed by two-dimensional gel electrophoresis (2D-gel), followed by mass spectrometry. The counting of gel spots indicated at least 800 proteins were present in these tissue, but due to technological restraints of the times, less than 100 were identified.^12^ We report herein the methodology and preliminary results of our attempts to create the first draft map of the *Cannabis* proteome. Proteins were extracted from plant tissue from stems and leaves of plants as well as from medical flower products from *C.sativa* and *C.indica* strains with well characterized cannabinoid profiles. Extracted proteins were digested, separated by ultrahigh pressure liquid chromatography and analyzed by ultrahigh resolution tandem mass spectrometry. Data assembly on the high- resolution spectra has been completed, resulting in the annotation of over 17,000 potential protein coding regions.

Traditional proteomics workflows rely on the existence of annotated theoretical proteins FASTA files derived from annotated genomes. Unfortunately, no annotated genome exists for any *Cannabis* plants. Data assembly on the high- resolution spectra has been completed, resulting in the annotation of over 17,000 potential protein coding regions lacking identifications. We have developed a pipeline by which any material with both “next generation” sequencing and shotgun proteomics data may be used to generate theoretical protein FASTA files directly, thus circumventing the need for annotated genomes entirely. The output of this pipeline is the most comprehensive protein FASTA for *Cannabis* constructed to date, consisting of 13,850 nonredundant sequences with putative annotations. With the creation of this large species-specific database we can now utilize traditional proteomics tools for the identification and quantification of proteins from these plants. Furthermore, we have identified diverse chemical modifications on proteins central to metabolism that appear linked to terpene and cannabinoid production by the plant. Correlating proteomics measurements with phenotypic data such as chemical profiles will provide a valuable resource for producers and concerned consumers and is a critical next step in the advancement of the medical applications of *Cannabis*.

To disseminate both the unprocessed and processed data we have established the website www.CannabisDraftMap.Org which features an interactive database of all progress on both this project as well as our progress on profiling the metabolomic pathways in these plants. **Table 1** is an overview of the current status of this project at the date of this submission.

**Table 1.** An overview of the progress to date.

|  |  |
| --- | --- |
| **Protein Sequenced** | **17,269** |
| **Protein Annotated** | 13,929 |
| **Proteins with homologous 3D structures** | 964 |
| **Acetylation sites Mapped** | 584 |
| **MS/MS Spectra Acquired** | 1.40E+07 |
| **MS/MS Spectra Searched** | 2.40E+06 |
| **MS/MS Spectra with Evidence of Glycosylation** | 3.50E+05 |
| **Skyline Spectral Library** | 43,612 annotated spectra |
| **Gene Coding Regions Annotated** | 13,850 |
| **Small Molecule Features Isolated** | 1,050 |
| **Small Molecules Identified** | 535 |

## Materials and Methods

### Samples

A table of samples analyzed to date are described in **Supplemental Table 1**. All samples were obtained by Think20Labs under the guidelines of the MMCC regulations in accordance with a temporary license granted under COMAR 10.62.33.^13^ A recent study described the optimization of digestion conditions for the proteomic analysis of *Cannabis* flowers and performed similar experiments as the ones described here. Vendor instrument files are available at the MASSIVE data repository as MSV00083191. MGF files for all data described in this study may be downloaded from www.CannabisDraftMap.org

### Sample Preparation

Multiple variations on protein extraction and digestion were tested, based on the highest percent recovery of peptides per milligram of starting plant material by use of an absorbance assay for tryptic peptides (Pierce) (data not shown). The final sample method was based on the filter-aided sample preparation method (FASP).^14^ Briefly, 1mg of fresh plant material was flash frozen at −80°C for 20 minutes. The cell walls were disrupted by immediately removing the frozen material and blunt physical concussion. The material was then dissolved in a solution of 150 μL of 5% SDS and 0.2%DTT and heated at 95°C for 10 minutes to reduce and linearize proteins and reduced to room temperature on ice. 150 μL of 8M urea/50mM TrisHCl was added to the mixture. Detergent removal, protein cysteine alkylation and sample cleanup for digestion was performed according to the FASP protocol. All reagents were obtained from Expedeon BioSciences. Proteins were digested for 16 hours at room temperature. Digested peptides were released by centrifuging the FASP chamber at 13,000 x g for 10 minutes with peptides eluting into a new 1.5 mL centrifuge tube. An additional 75 μL of the digestion solution was added and the elution was repeated. The peptides were dried by vacuum centrifugation (SpeedVac). Peptides were resuspended in 20μL of 0.1% trifluoroacetic acid for either desalting or for high pH reversed phase fractionation. Peptides were quantified by absorbance using a peptide specific kit (Pierce).

### Peptide Fractionation

Alkylated peptide aliquots of approximately 50 micrograms were combined from each sample into a shared pool for offline fractionation and library generation. Due to sample availability constraints, two batches containing peptides from 6 separate samples were fractionated separately. High resolution fractionation followed a recent protocol^7^, with the exception that an Accela 1250 pump (Thermo) was utilized for gradient delivery and fractions were concatenated following collection. The concatenation was performed as described previously.^15^

### LC-Mass Spectrometry Analysis

For initial analysis all fractionated and single shot samples were ran identically on a nanoESI-Q Exactive HF-X system. Briefly, 4 μg of peptides were loaded into a 4cm trap column and eluted with an optimized gradient on a 100 cm monolithic 75μm column. Eluting peptide masses were acquired at 120,000 resolution followed by the fragmentation of the most abundant eluting peptides with HCD fragmentation at 27eV. Fragmented peptides were acquired at 15,000. Although this system is capable of higher scan speed, a higher resolution MS/MS was utilized in order to obtain more confident identification and localization of PTMs. The top 15 most abundant ions were selected for fragmentation. Each single shot sample was injected twice, once with a 30 ms ion injection time, and again with a 150 ms ion accumulation time for each MS/MS scan. Dynamic exclusion was utilized allowing each ion to be fragmented once, any ion within 5ppm of the matched ion was excluded from fragmentation for 60 seconds, or approximately 2.2x the peak width.

### Peptide and Protein Identification

An overview of the data processing pipeline and all input is demonstrated in **Figure 1**. To date, no full annotated protein FASTA exists for any *Cannabis* species. Classical proteomics workflows require a reference theoretical protein database from which to construct matches from MS1 and MS/MS spectral data. In lieu of this we utilized 2 sources of information for identifying MS/MS spectra. As less than 600 annotated sequences for *Cannabis* exist in the UniProt library, a custom UniProt/SwissProt database consisting of every manually annotated sequence from green plants was used. In conjunction with this the three highest quality genome sequences available in the literature were subjected to 6-frame translation in house using the MaxQuant v1.6.3.3. software suite to create theoretical protein sequences that accurately match the material being analyzed. These files will be referred to herein as our proteogenomic FASTA. For initial analysis, all data processing was performed in Proteome Discoverer 2.2 (PD) (Thermo Fisher) using the SequestHT, Percolator and Minora algorithms. The proteogenomic FASTA was crudely reduced during database import in PD according to manufacturer default settings. SequestHT and Percolator generate identity and confidence scored peptide spectral matches (PSMs). Multiple consensus workflows were used within PD to assemble the PSMs into peptide groups, protein database matches, and finally non-redundant proteins groups using the principle of strict parsimony as defined by the vendor software defaults. All settings utilized in the data processing to the generation of PSMs and the Consensus steps that reduce these matches to protein group identifications are described in **Supplemental Table 2**.

**Figure 1.**
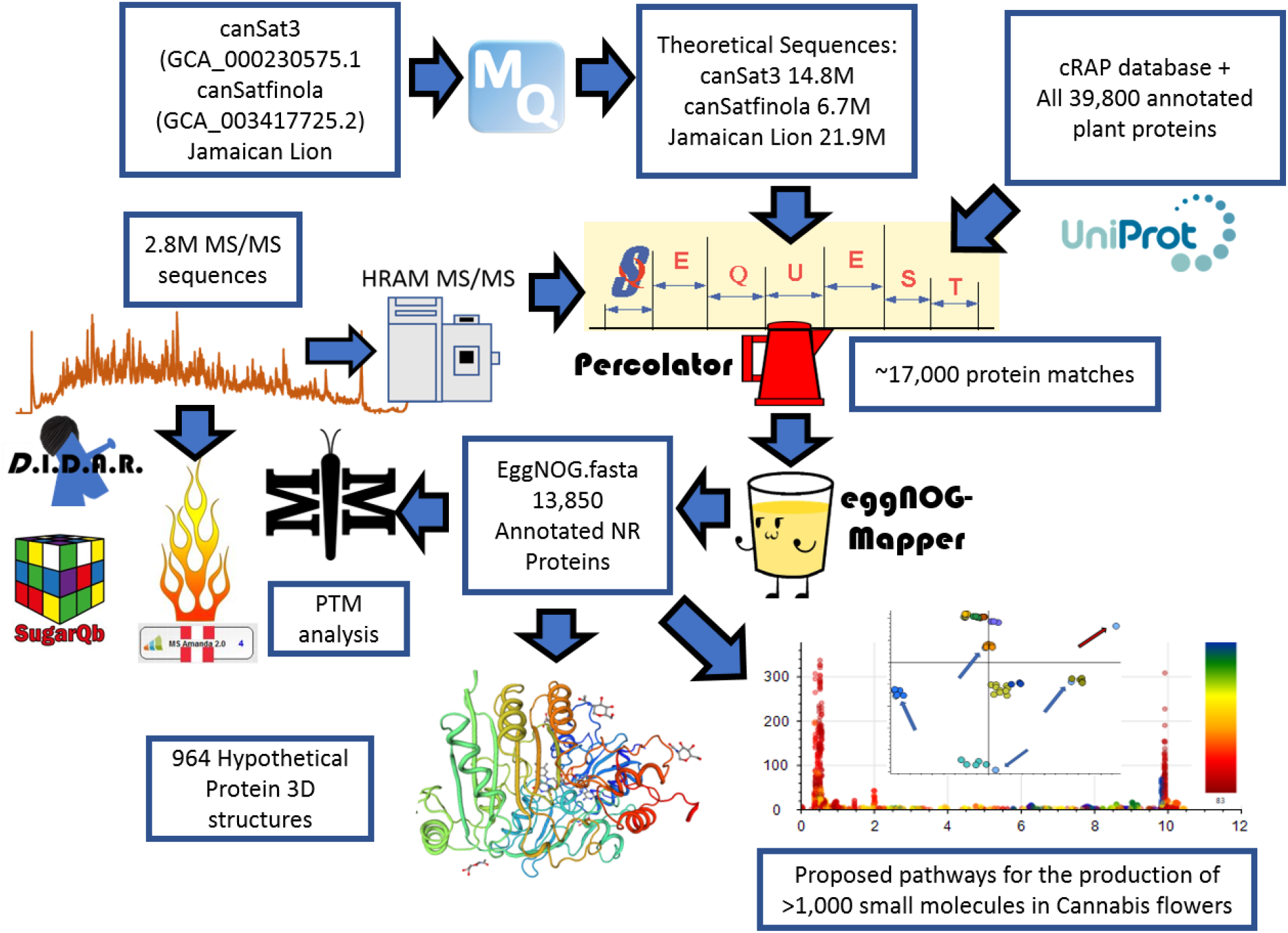
A cartoon describing the bioinformatic pipelines leading to the current results and resources present on www.CannabisDraftMap.org.

### Generation of the eggNOG annotated protein FASTA

The 6-frame translated database contained approximately 43.4 million potential proteins sequences. Of these, 86,944 had at least one PSM uniquely matched to the proteogenomic FASTA or the UniProt Viridiplantae FASTA file, 58,309 were found to be non-redundant by sequence. All candidate protein sequences with at least one PSM were utilized to generate the V1.0 *Cannabis* annotated FASTA in the following manner. All proposed proteins sequences were exported from Proteome Discoverer 2.2 in FASTA format. To reduce and annotate this file, the eggNOG-mapper program^16^ was utilized with default server parameters with annotations utilizing any orthologs from the Viridiplantae database online as of the date of utilization (07/04/2019). The returned files contained 30,988 putative functional annotations. 5,735 entries had significant homology under the default server parameters for assignment to a specific gene. Using these settings, 796 sequences could not be assigned a significant match to the database for functional annotation. Further investigation will be necessary to determine if these proteins are artifacts of data processing or sequences uniquely present in these plants. The remaining 24,457 sequences were assigned a protein accession and functional annotation based on sequence. In order to preserve a unified format required for Proteome Discoverer, the gene name was replaced with the phrase “gene not found” when the best annotation was by protein function. The eggNOG annotation file and resulting FASTA were merged using an in-house generated script. The final annotated FASTA was compiled with the FASTA database utilities tool in Proteome Discoverer 2.2 and the compiled database was uploaded into the program, resulting in a final annotated database with 13,850 non-redundant annotated protein sequences.

### Spectral Library Generation

The high-resolution MS/MS spectra were searched with PD 2.1 using the SeQuestHT algorithm and eggNOG FASTA file using the same settings as described above and in **Supplemental Table 2**. The resulting

.pdresult file was imported into the Skyline^17^ 64-bit environment (version 4.1.0.1869) and converted to a spectral library according to the default parameters for Orbitrap high resolution MS/MS spectra.^18^ The output spectral library can facilitate both targeted and Data Independent Analysis of plant proteins and is available for download at www.CannabisDraftMap.org

### Chromosome alignment

A recent re-analysis of the CanSat3 genome^19^ aligned the sequences into ten separate chromosome files.^20^ The Protein Marker node in Proteome Discoverer was used in 4 rounds of reprocessing of the consensus workflow to develop a metric of the number of identified protein entries in this study that are products of each chromosome. Four rounds were necessary due to a limitation in the software that allows a maximum of 3 separate FASTA sequences to be used for output marking. Reiterations of this analysis were repeated to ensure that the chromosomes grouped in each re-analysis was an independent variable and did not affect localization output (data not shown). Both protein data, representing potential redundancies and unique protein groups were obtained. Using an exact match approach, 3,421 proteins could be confidently mapped to one or more chromosomes. Cannabis has 10 pairs of chromosomes. Using this approach alignment is only possible to matching pair, not individual chromosome. The results are plotted in **Figure 2**.

**Figure 2.**
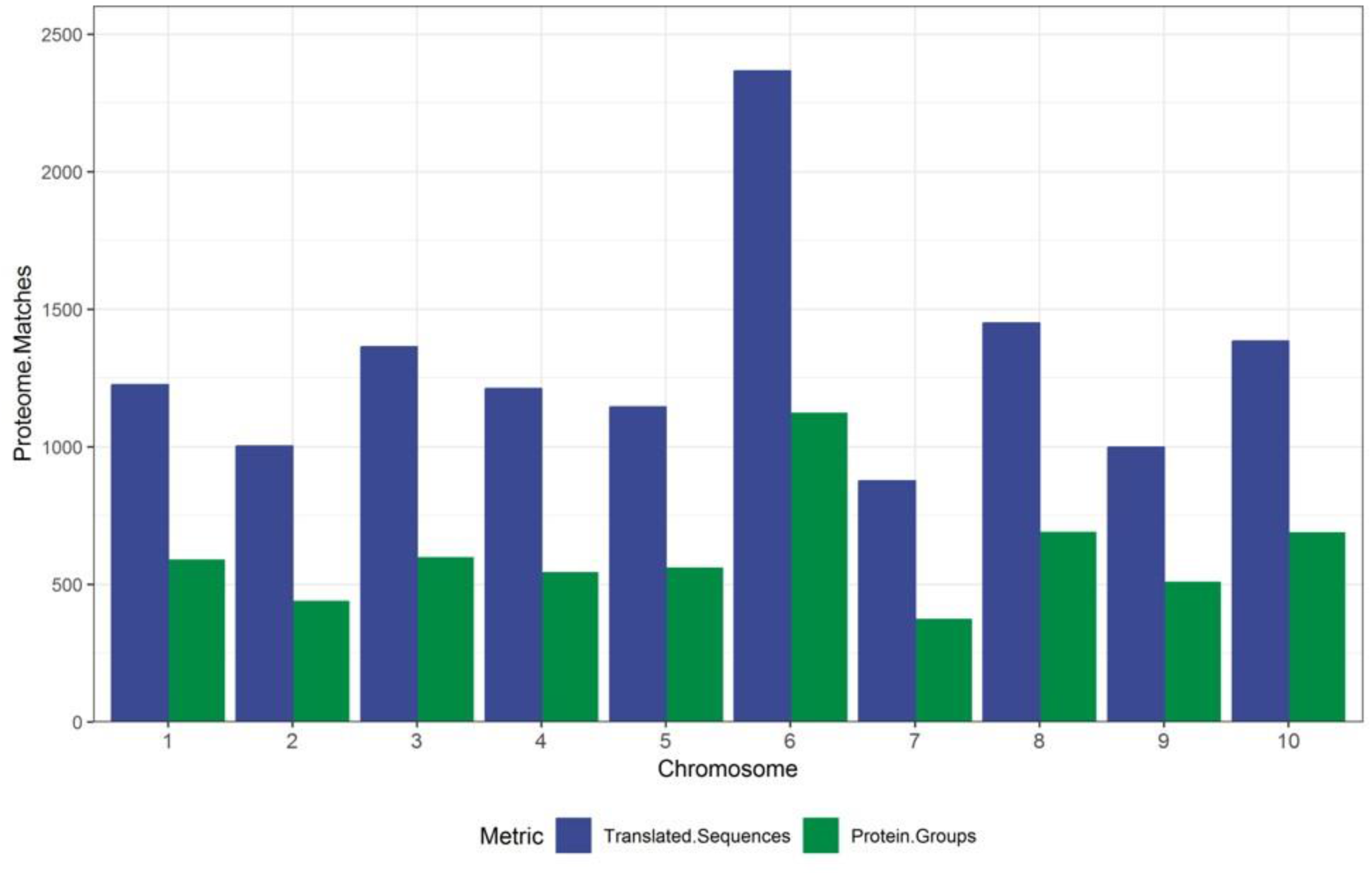
A graph demonstrating the alignment of the proteomic presented here against the CanSat3 chromosome assemblies.

### Identification and validation of post-translational modifications (PTMs)

The MetaMorpheus software package v0.0.301 (MM) was used for the indiscriminate identification of post translational modifications. The unannotated FASTA file was used for MM analysis using the default workflows for Recalibration, GPTMD, Search and Post Processing^21,22^ using default parameters. A resulting output file is **Supplemental Table 4**. To further confirm and visualize the presence of the most abundant PTMs, lysine acetylation and serine/threonine phosphorylation, IMP-PD 2.1 (pd-nodes.org) was utilized. The workflow consisted of MSAmanda 2.0 operating with 5ppm MS1 and 15ppm MS/MS tolerance. The search was performed with eggNOGFASTA v.1.0, the Viridiplantae UniProt FASTA and cRAP. Static modification of carbamidomethylation of cysteine, with dynamic modifications of methionine oxidation, lysine acetylation and phosphorylation of serine and threonine were all enabled. The ptmRS algorithm^23^ was used for confidence of site localization. All consensus workflow settings matched those described for the SeQuest searches, with the exception that the localization of any PTM with 50% likelihood or greater was allowed for visualization. The pdResult output file was visualized in MS2Go v1.4.7 (www.pd-nodes.org) according to default parameters. Files were filtered at the PSM level for lysine acetylation and phosphorylation of serine/threonine, respectively.

To search for the presence of glycosylated peptides, the SugarQb program^24,25^ was used in Proteome Discoverer 2.1 using the default workflow as described in the instruction manual on pd-nodes.org. For this analysis only one fractionated file set was searched, containing the combined flower samples 7-12 that were high pH fractionated as described above. These samples were analyzed in an identical manner to the samples above, with the exception that an MS/MS fill time of 150ms was used for the accumulation of the 15,000 resolution fragment ions. A first mass of 100 m/z was set in order to observe the presence of diagnostic ions necessary for the SugarQB and Diagnostic Ion Data Analysis Reduction (D.I.D.A.R.) software. All settings and potential glycan structures used in the SugarQB workflow are provided as **Supplemental Tables 7** and **8**, respectively.

### 3D Protein Library Construction and Similarity Scoring

EggNOG-mapper derived similarity accession numbers were combined via in-house script with the generic URL output extension for Swiss-Model library functions.^26,27^ A spreadsheet containing the 965 proteins with Swiss Model entries is provided as **Supplemental Table 10**.

### Correlation Analysis of Small Molecules and Proteins

We have recently described the identification of approximately 1,000 distinct small molecule features present in extractions from mature cannabis flowers.^28^ Six samples used in this previous study were also prepared for single shot LC-MS analysis using a variant of the IonStar methodology^29^ as described above. These individual files were processed in PD 2.2 using SeQuest searched against the eggNOG FATA and utilizing the Minora algorithm for relative label free quantification according to manufacturer default settings. Quantification values were derived from pairwise analysis at the peptide group level. Abundance ratios were averaged in when two or more protein groups were observed. The resulting file contained 3661 protein groups and is provided as **Supplemental Table 11**.

A correlation analysis was then performed on the small molecule features and the label free quantification results utilizing Python (v3.7.3) along with the Pandas (v0.25.1) and Scipy (v.1.3.1) packages (https://github.com/jenkinsc11/probocor). For each small molecule that was identified, the changing areas of the features between samples were directly compared to every identified protein’s changing calculated abundances. Pearson and Spearman correlation values were calculated along with their respective p-values. If either correlation analysis had the arbitrary p-value cut-off of less than or equal to 0.05, the small molecule – protein quantitation change between samples was flagged as having a possibly statistically significant correlation and compiled into a list for further investigation.

To verify the power of this approach, several molecules were manually evaluated by extracting the molecules of interest to the proteins with the strongest Pearson Correlations. A description of this process is demonstrated in **Figure 5**. The molecule 11-OH-THC was found to most strongly correlate to the protein with eggNOG accession 981085.XP_010108776.1 with a Pearson Correlation of 0.9997 and a corresponding p-Value of 1.54 ×10^−7^.

### Graph generation

The UpSetR package was used for the comparison of proteome to genome sequencing files using both the webhosted ShinyApp (https://gehlenborglab.shinyapps.io/upsetr/) as well as the full package within RStudio 1.0.143. Supplemental figures were generated in the PNNL Venn Diagram 1.5.5 tool as well as with GGPlot2^30^ within RStudio.

## Results and Conclusion

### Peptide and protein identifications

The near-complete lack of annotated genetic sequences and theoretical protein sequences is a considerable challenge for traditional proteomics workflows which rely heavily on these resources for spectral matching. Using the custom proteogenomic workflow described here, we were able to identify a small percentage of the first 2.5 million MS/MS spectra obtained and match those to a compiled and in-house generated theoretical protein sequence database of greater than 41 million entries. This pipeline resulted in 135,845 peptide spectral matches, or approximately 5.4% identification rate.

A primary focus of future work will be the refinement and annotation of the currently existing Cannabis genomes available. Recent work has described the improvement and correction of genome annotations using high resolution mass spectrometry.^31^ While this is beyond the scope of this study, we can develop metrics related to the quality of match of genomic data using high coverage proteomics. An UpSetR graph^32^ is shown in **Figure 3** that shows the unique protein identifications and matches to the various genomic databases both unique and conserved. To further illustrate the importance of an annotated Cannabis protein database, of the protein groups identified by the initial analysis, only 1,838 representing less than 10% had sequence homology suitable to be matched against the UniProt/SwissProt database, despite the fact a database containing the protein sequences of all green plants was utilized in this work. A summary of these results is available as **Supplemental Table 2**.

**Figure 3.**
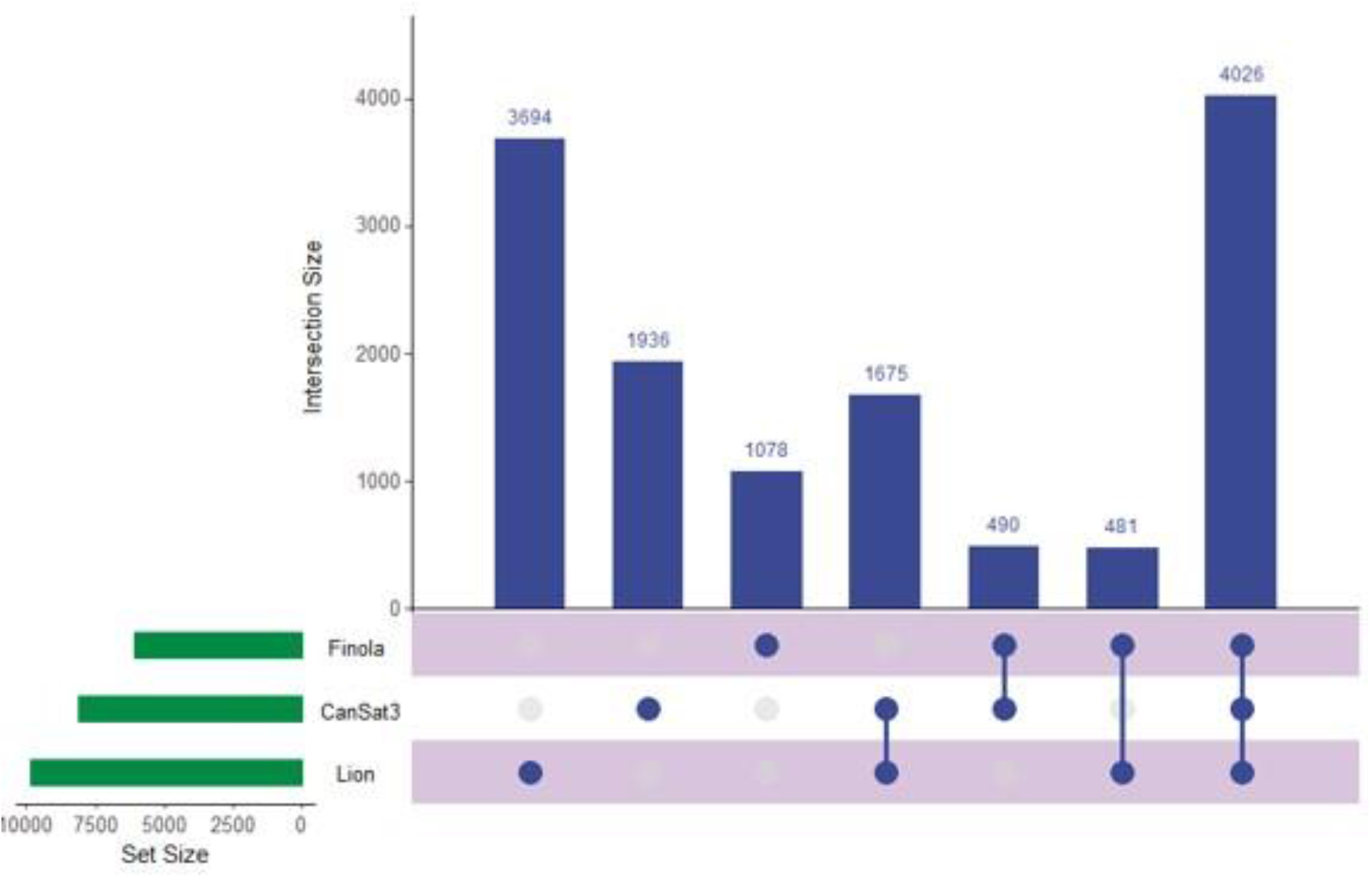
A graph demonstrating the alignment of the proteomic data presented here to previous genome sequencing projects

Despite these challenges we have built considerably on the existing knowledge of the *Cannabis* proteome and have generated a relatively complete picture of the protein composition of the plants we have analyzed to date. Genomic analysis has identified approximately 25,000 protein coding regions, a number comparable to that of humans in a Cannabis chromosome assembly^33^. While exact numbers are still currently debated, no study to our knowledge as identified more 20,000 or more unique protein groups in humans.^34^

### Post-translational Modification Identification and Analysis

The importance of protein post-translational modifications in Cannabis are, to our knowledge, completely unknown, outside of results suggesting glycosylation on the THCA synthase protein.^35^ Current strategies for identifying PTMs from shotgun proteomic data require the addition of dynamic modification. Each single dynamic modification results in a doubling of the number of theoretical peptides and due to the presence of multiple modifications sites per proteins, indiscriminate searching of PTMs results in exponential increase in both the search space and required computational power to complete data processing.^36^ To address these issues we generated a new FASTA database that contained only the 17,269 proteins identified in our SeQuest and Percolator searches of all high resolution files. Using this newly reduced database of proteins that appear to be present and the complete theoretical sequences from these entries extracted from our original FASTAs, we can search these identified proteins for PTMs. For this analysis we chose to employ the recently described MetaMorpheus (MM) software. MM performs a tiered search strategy that is reliant on the recalibration of MS spectra and the GPTMD algorithm.^37^ This next generation search engine is capable of searching for, identifying and quantifying hundreds of unknown post-translational modifications with annotated databases on standard desktop computers.^21,22^ **Supplemental Table 4** contains a summary of these results. MM identified 26,477 unique peptides and 6,111 PTMs in these files alone. The most common identified modification was methionine oxidation, which is often a product of the sample preparation process. Lysine acetylation appeared in a high number of PSMs and phosphorylation of serine and threonine were also observed. To further confirm, localize and visualize these potential PTMs, the fractions from 6 flower samples and the fractions from male plant stems and leaves, respectively, were combined and searched with MSAmanda 2.0 and the ptmRS software.

The MSAmanda search engine was specifically designed for high resolution accurate MS/MS spectra and has been demonstrated to be a particularly powerful in the confident identification of PTMs. The ptmRS algorithm provides scoring of the probability of site localization of PTMs within the peptide chain and is most useful when multiple amino acid residues may host this chemical modification. MSAmanda 2.0 is a recent iteration of the software which allows far more practical sequencing speeds on standard desktop computers. Data visualization was performed using the recently described MS2Go software (www.pd-nodes.org). The complete MS2Go output is available for download at www.CannabisDraftMap.org under the Full Data Sets page.

A total of 584 proteins were identified that possessed at least one lysine acetylation within their protein sequence. MS2Go output displaying these sequences is presented as **Supplemental Table 5**. The global analysis of lysine acetylation by shotgun proteomics has only been recently described but the application of this technology has demonstrated the presence of this PTM in diverse proteins across many species.^38^ Lysine acetylation has recently been described as a key modulator of the model organism *Arabidopsis thaliana*.^39^ The design of this analysis presented here does not, by design, allow quantification. However, we were interested to find that the over 90% of the observed acetylation sites were unique to mature flowers and appeared entirely absent in other plant materials. Furthermore, lysine acetylation sites were observed in proteins involved in the production of compounds of commercial interest. **Supplemental Figure 3** is a visualization of the sequence coverage and acetylation sites of the THCA synthase protein (THCAS). Three acetylated peptides were found in the TPS1 gene product which is involved the final stages of the synthesis of limonene, **Figure 4** is a depiction of one peptide from this protein of interest. In addition to peptide sequence coverage, HCD fragmentation of lysine acetylated peptides typically results in the production of a diagnostic fragment ion of 126.0913.^40^ All peptides demonstrated as acetylated in THCAS and TPS1 were further supported by manual validation of the presence of this diagnostic ion, although not within the scale of the image for the spectra chosen for TPS1.

While of questionable medicinal value^41^ the characteristic limonene fragrance is of specific commercial value and the state of CA requires the quantification of this terpene in all Cannabis products. Further investigation is underway to determine the relative importance of this PTM in the flower and the production of secondary metabolites of commercial and medical value.

A previous study suggested the possible existence of glycosylation on THCAS at up to four sites. This conclusion was based on a the presence of an unexpected gel shift during electrophoretic migration of purified protein and the presence of structural domains compatible with glycosylation.^20^ Given the relatively high abundance of this protein, we found it surprising that few glycopeptides were predicted by the MetaMorpheus analysis. To further investigate we used the recently described SugarQb algorithm and an in-house developed software called D.I.D.A.R to investigate the highly fractionated samples from the combined protein extracts of 6 medicinal flowers. A slightly modified LCMS instrument method was utilized on the instrument to allow a greater fill time and to allow lower mass MS/MS acquisition necessary for picking up diagnostic fragment ions. SugarQb can identify glycopeptides from tandem mass spectra by first searching for fragments indicative of the potential modification and iteratively searching those spectra with glycan libraries added into the workflow. We utilized a library obtained from the IMP proteomics facility that considered the possibility of up to 1,575 unique glycan chain modifications, in addition to the single modification of N-acetylglucosamine (GlcNaC) on all amino acids. Although 174 glycosylation sites were identified, none were localized to the proteins involved in the synthesis of cannabinoids. The full list of glycosylated proteins is provided as **Supplemental Table 8**.

**Figure 4.**
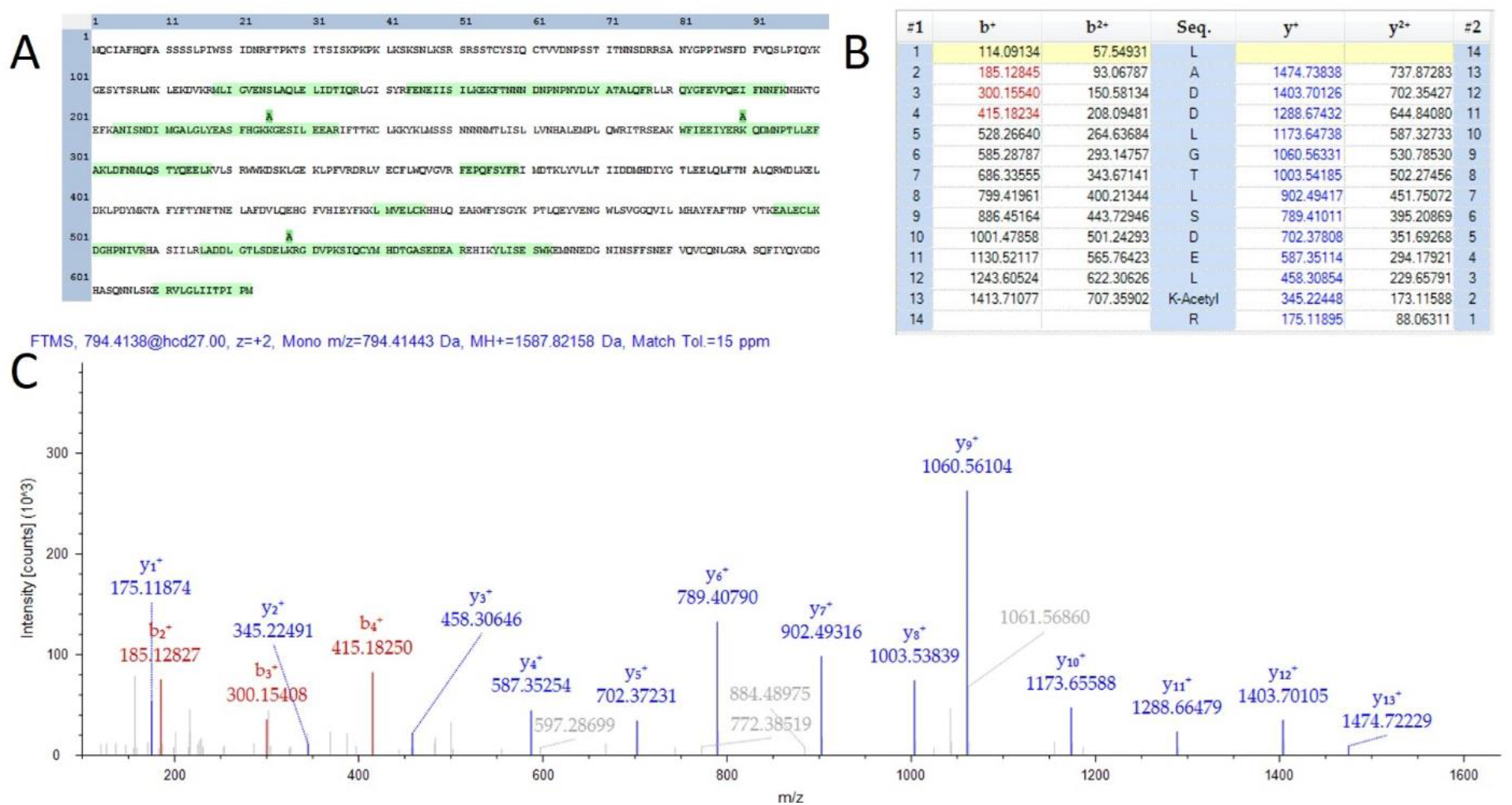
Evidence of acetylation on TPS1. **A)** A sequence map demonstrating 3 observed lysine acetylation modifications. **B)** A fragment map showing 100% sequence coverage for one acetylation site. C) MS/MS spectra matching the fragment map

**Figure 5.**
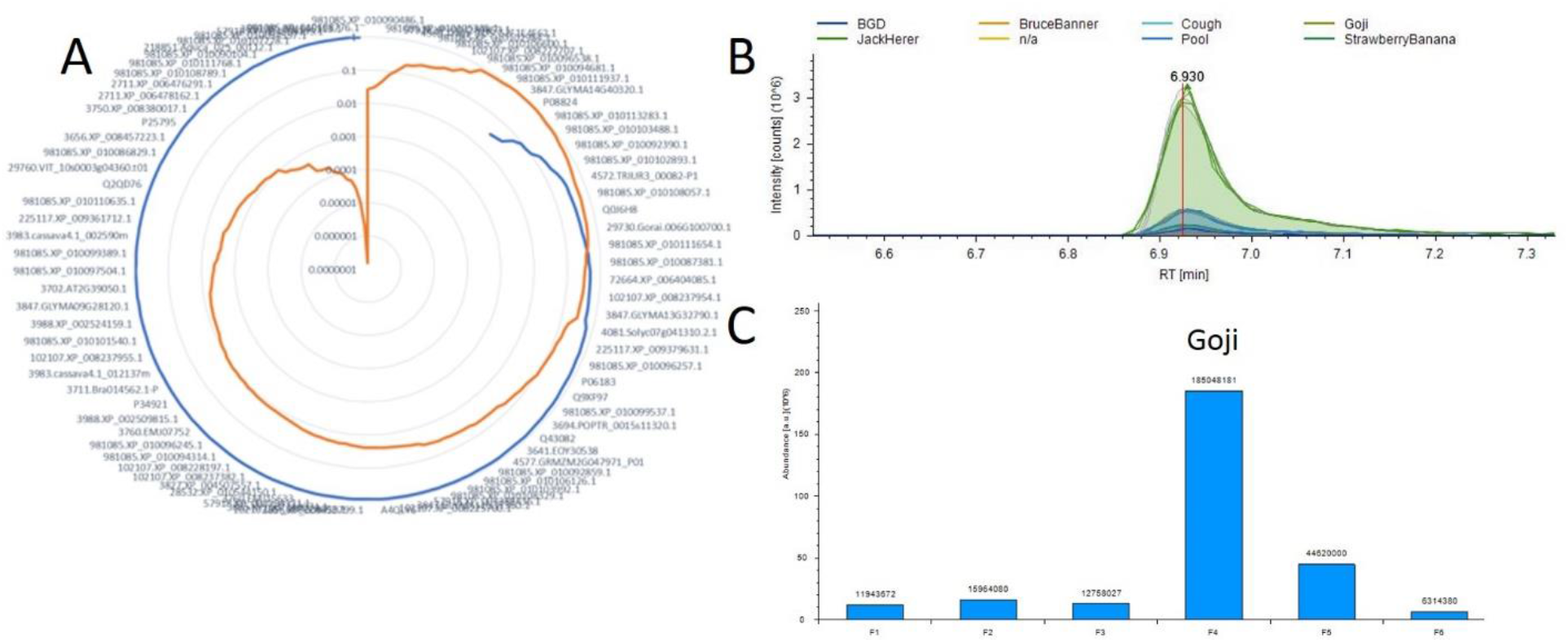
Example correlation analysis plot **A)** Radar diagram for 11-OH-THC where the blue line represents all positive Pearson Correlation and orange is the p-value for each measurement. **B)** A plot overlaying the 11-OH-THC metabolite peaks and technical replicates. **C)** A plot of the protein from A demonstrating the highest correlation with this metabolite.

We have recently described the Reporter Ion Data Analysis Reduction (R.I.D.A.R.) software.^36^ This software has the capability of removing spectra that are not quantitatively interesting to the end user from large shotgun proteomics datasets prior to database searching steps. By reducing the data in this manner, we can lower the processing time of large cohort studies to be manageable for processing on a standard consumer level computer equipment. Diagnostic Ion Data Analysis Reduction (D.I.D.A.R.) is an extension of this logic and will be described in depth elsewhere. The python script allows an end user to remove spectra that contain only those with fragment ions of interest.^40^ We used the D.I.D.A.R. script to create a file containing only spectra with the diagnostic fragment ions. The diagnostic ions as well as the number of MS/MS spectra containing these markers is provided as Supplemental Table 13. Surprisingly, over 3 ×10^5^ MS/MS spectra returned as positive for glycan specific fragment ions. To verify the validity of this output, we used the Xcalibur software to display an extracted ion chromatogram plotting only the signal of ions present in MS/MS spectra with an m/z within 5ppm of the HexNaC diagnostic fragment ion. This visualization is shown in **Supplemental Figure 4**. As none of these ions appeared to match our SugarQb library, further investigation is underway to determine if Cannabis plants use a currently unknown glycosylation mechanism.

### Three-Dimensional Protein Structures

Protein 3D models can allow a more thorough understanding of protein-protein interactions, particularly in metabolic processes.^42^ At the date of this writing the Swiss-Model database contained only two experimentally determined crystal structures for proteins from Cannabis. As a demonstration of a potential application of the Cannabis proteome draft map and eggNOG derived database, we used the eggNOG derived accessions to generate a list of homologous proteins with existing SwissModel entries form green plants. The resulting output is provided as **Supplemental Table 10** and contains 964 hyperlinks to existing models. Further work will be necessary to determine the strength of these 3D model determinations from the linear homologies shown here. The modeling of Cannabis proteins by 3D homology is beyond the scope of this work but may provide further insight into the inner working of the biochemistry of these plants.

### Utilizing this resource toward the meta-analysis of previous studies

A recent study from Vincent *et al*., sought to develop an optimized protocol Cannabis proteomics.^43^ This study utilized a linear ion trap Orbitrap (LTQ Orbitrap Velos) system and nanoflow HPLC system ultimately similar system to those employed in this work. While the focus of the work was digestion efficiency, the number of proteins identified were low by comparison, with less than 200 total protein identifications. The authors point out that this is due to the small number of annotated proteins present in the UniProt database.

We reanalyzed the instrument vendor files from this work using the eggNOG FASTA databases developed in this study. The complete output file is available in **Supplemental Table 9**. Using our UniProt derived Viridiplantae database, we find that this work matches 330 confident protein groups (in sum). By adding the eggNOG .fasta database to this identical analysis we obtain a total of 2,026 high confidence unique protein groups from the combination of all deposited instrument vendor files.

### Correlation Analysis Between Small Molecules and Proteins

Correlation analysis in proteomics has been shown to be a powerful tool in the identification of cooperating proteins in biological processes.^44^ We have recently described the identification and quantification of over 1,000 small molecule features in medicinal products. While the pathways leading to the production of the major cannabinoids has been the focus of much study, little is known regarding the production of other central and secondary metabolites. The lists of small molecules and proteins with quantification values were combined resulting in correlation scores and significance values linking all metabolites to confidently identified and quantified proteins.

Using this tool as a starting point we hope to map all metabolic pathways in *Cannabis* plants and to identify both new molecules of interest and the proteins responsible for their production. Current work is focused on the acquisition of more and varied samples to further develop this resource.

## Conclusions

We have performed the first comprehensive proteomic analysis of *Cannabis* plants toward our goal of developing a multi-omics biochemical map of these plants. From the samples analyzed to date and described herein, we have identified a possible 17,269 unique proteins. To circumvent the lack of a comprehensive annotated genome for any *Cannabis* species, we have created the first annotated protein FASTA database for these plants, comprising nearly 14,000 proteins annotated to gene identities when possible, and grouped by putative function if not. The possession of the FASTA database makes proteomic analyses possible with existing software. We have found that *Cannabis* plants possess numerous post-translationally modified proteins, namely lysine acetylation sites and phosphorylation of threonine and serine, as well as evidence of extensive glycosylation. The analysis of the role of phosphorylation in *Cannabis* plants is outside the scope of the currently study, as thorough analysis requires chemical enrichment, lysine acetylation appears to be involved in the production of Cannabis molecules of commercial and medical interest. The use a of the new DIDAR algorithm suggests the presence of multiple post translational modifications, with glycosylation appearing to play a major role in plant biochemistry. To facilitate further study of these plants, we have made our FASTA database, annotated spectra and spectral libraries publicly available with the release of this manuscript, along with other resources at www.CannabisDraftMap.org.

### Future goals

We aim to identify the function of PTMs in these plants, specifically how these modifications correlate to the production of secondary metabolites. Multiple alternative algorithms and approaches may be used to further refine, improve and annotate all resources described in this study and investigation into these approaches are currently underway. Furthermore, it is our belief that big data is only useful if it is made available to the largest possible audiences. We will continue to work on clarifying our results and making these available to the wider community through improved software and interfaces, with a specific focus on improving and expanding on the spectral libraries available here to enable robust targeted and data independent acquisition analysis of these plants. All results described in this study and updated results will be made freely available at www.CannabisDraftMap.org

## Acknowledgements

We would like to thank Dr. Miranda Darby of Hood College Bioinformatics for helpful conversations on genetics and sequence alignment that clarified our path toward the construction of the eggNOG database.

## Supplemental Table Key

Supplemental Table 1: Samples Analyzed

Supplemental Table 2: Proteome Discoverer search settings Supplemental Table 3: UniProt Matches

Supplemental Table 4: MetaMorpheus_PSM_PTMs

Supplemental Table 5: Acetylated Peptides

Supplemental Table 6: SugarQB Settings

Supplemental Table 7: SugarQB Library

Supplemental Table 8: SugarQB output

Supplemental Table 9: Vincent *et al.*, Orbi Velos reanalyzed output

Supplemental Table 10: 3D Protein Structures

Supplemental Table 11: Full Correlation Analysis

Supplemental Table 12: 11-OH-THC Correlations

Supplemental Table 13: D.I.D.A.R Settings and Results

---

**Supplemental Figure 1:**
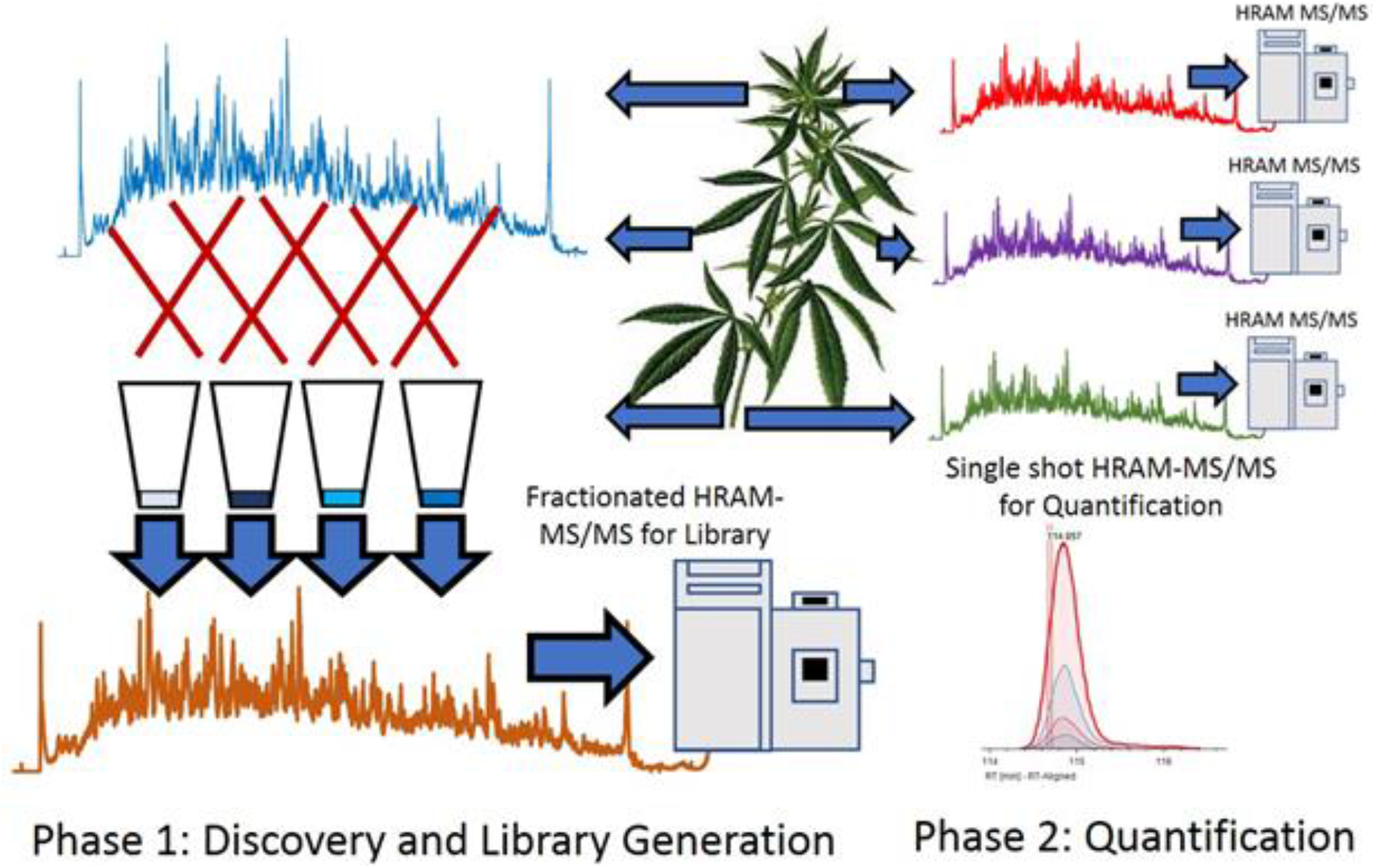
Library generation strategy Figure 1) An overview of the proteomics strategy. Combined material was heavily fractionated in order to develop library data. individual materials were separately analyzed by single-shot analysis for quantification.

**Supplemental Figure 2:**
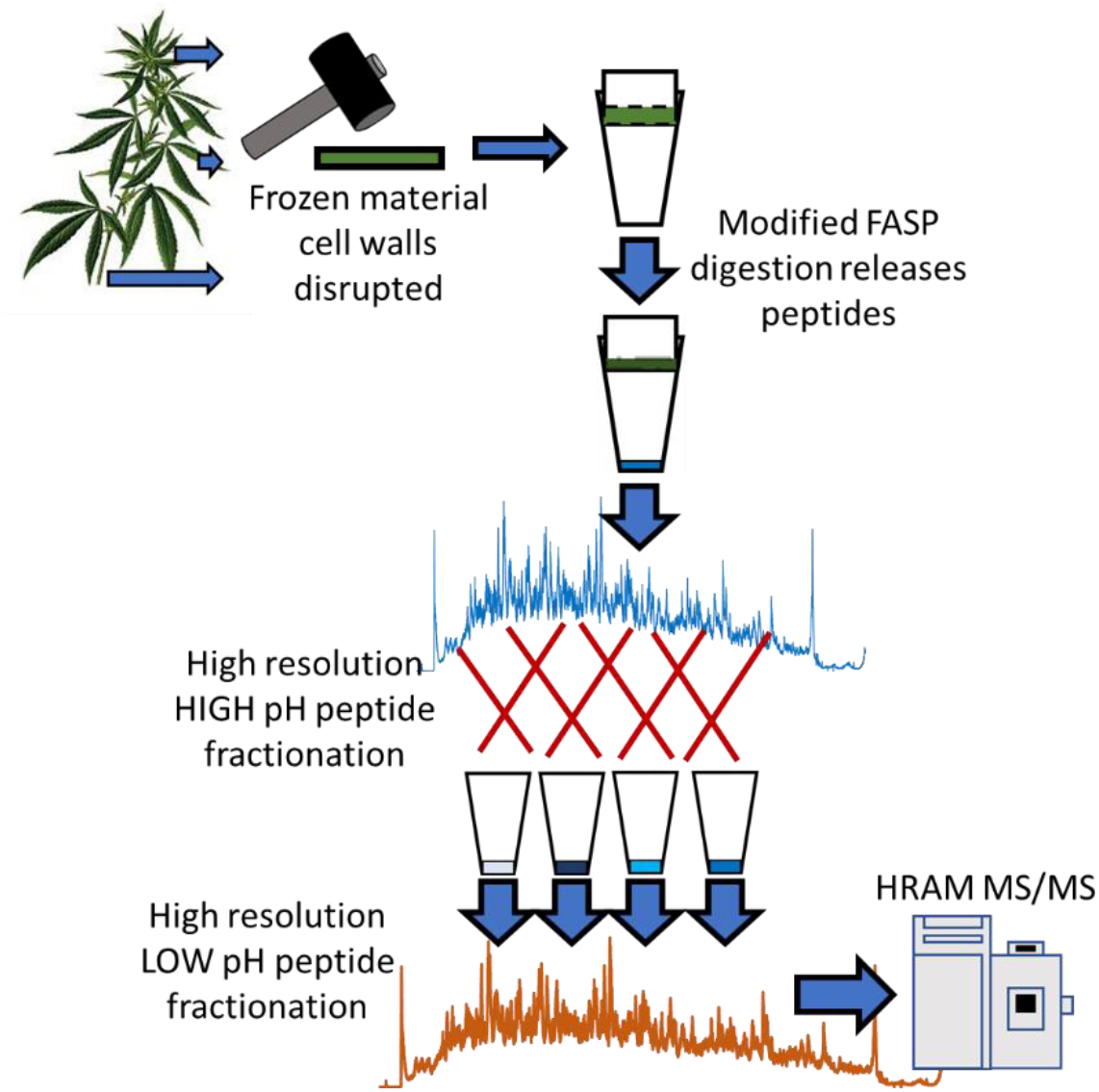
Protein extraction and digestion strategy cartoon:

**Supplemental Figure 3:**
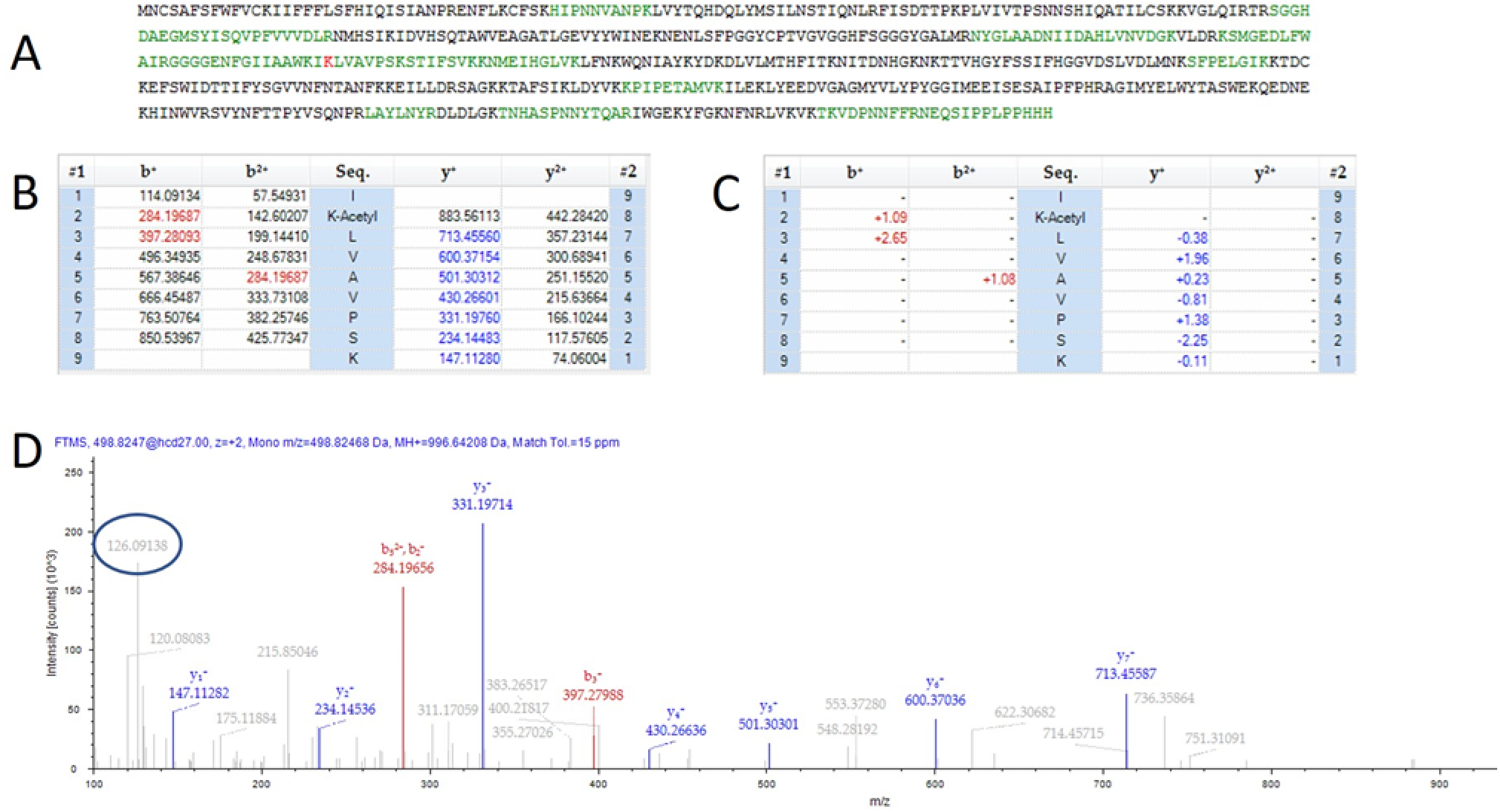
Possible THCAS Acetylation site: Figure 3A) The full theoretical sequence of THCAS with flagged lysine acetylation site in red B) The theoretical peptide fragment mao for the lysine acetylated peptide. C) The mass error of fragment ions identified in the MS/MS spectrum in panel D from theorectical. Values expressed in parts per million. D) The MS/MS spectrum identified as the lysine acetylated peptide lk(acetyl)LVAPSK. The circled fragment demonstrates a mass matching the persence of a lysine+acetyl diagnostic ion.

**Supplemental Figure 4:**
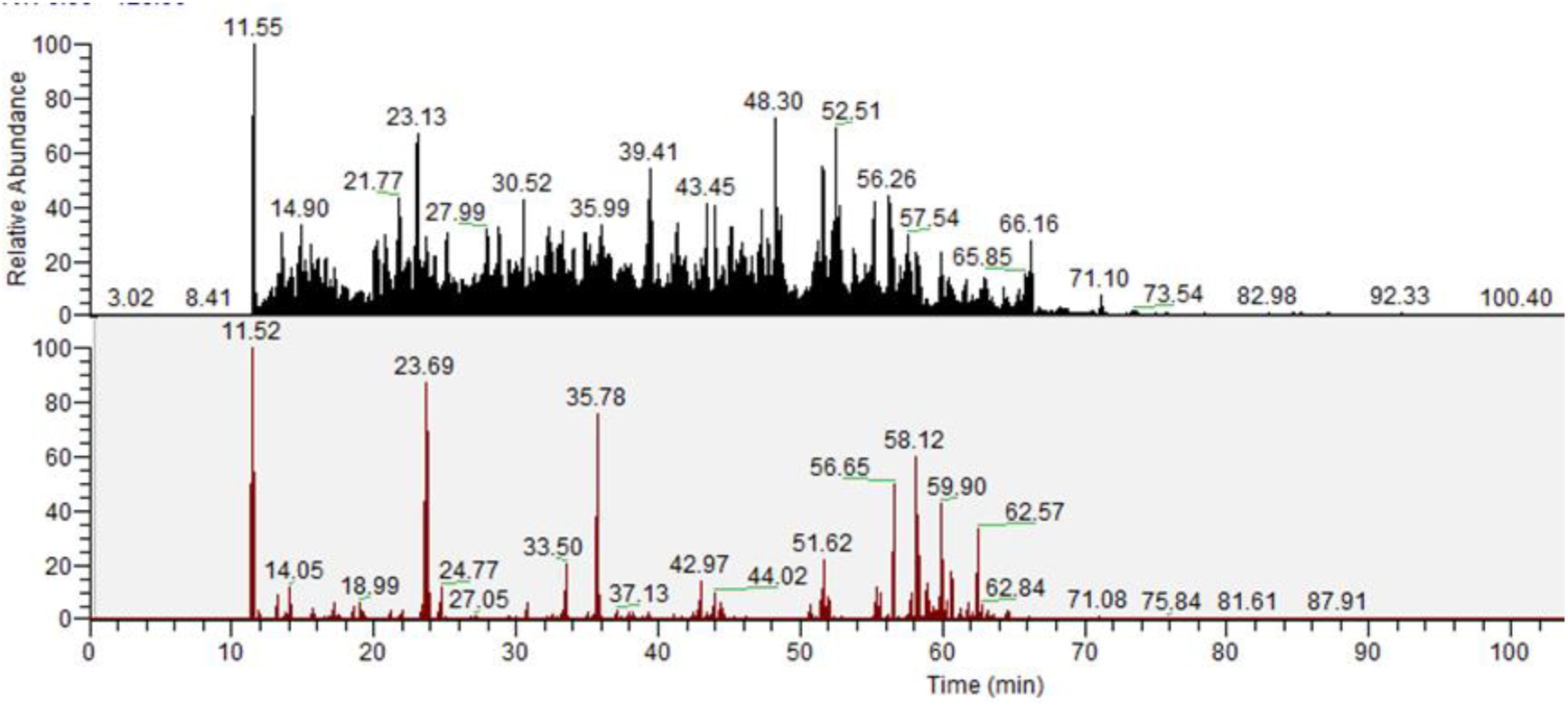
(Top pane) The total ion chromatogram from an arbitrarily chosen file from this study. (Bottom pane) The same chromatogram filtered to display only the signal of MS/MS produced ions within 5 ppm of the HexNaC diagnostic ion (204.0866)

